# Transcriptional changes consistent with altered neuronal differentiation, angiogenesis, and tumor plasticity induced in human subpallial telencephalic organoid-glioblastoma chimeras

**DOI:** 10.1101/2023.05.11.540229

**Authors:** Simone Chiola, Jingye Yang, H M Arif Ullah, Kandy Napan, Qiju Huang, Nicholas Gamboa, Osama Youssef, Howard Colman, Samuel H. Cheshier, Alex Shcheglovitov

## Abstract

Glioblastoma multiforme (GBM) is one of the most aggressive and therapy-resistant brain tumors prevalent in both adults and children. Despite extensive research to understand GBM pathology, it remains unclear how neural cells in the human brain interact with GBM cells to support their brain propagation and therapy resistance and whether GBM cells exert any influence on the properties of human neural cells. In this study, we co-culture human stem cell-derived subpallial telencephalic organoids with patient-derived proneural or mesenchymal GBM spheroids to investigate their reciprocal interactions. We show that both proneural and mesenchymal GBM spheroids readily fuse and propagate with human organoids, forming organoid-GBM chimeras, without the need for exogenous growth factors. GBM cells within the chimeras adapt by modulating gene expression profiles consistent with diminished proliferation, heightened hypoxia, increased angiogenesis, and proneural-to-mesenchymal transition in proneural GBM. Both proneural or mesenchymal GBMs also exert an impact on the properties of neural cells in the chimeras, leading to the suppression of neuronal genes and an upregulation expression of genes associated with hypoxia and angiogenesis. Collectively, this study identifies specific genes and molecular pathways that can be altered in GBM and neural cells by reciprocal interactions in a human developing brain-like environment for an increased understanding of GBM pathology and future therapy development.

## Introduction

Glioblastoma (GBM, WHO Grade IV IDH-wildtype diffuse astrocytic glioma) is one of the deadliest primary brain tumors affecting individuals of all ages (Brat et al. 2018; Ostrom et al. 2019; Louis et al. 2021). Although GBM has been extensively investigated in xenograft animal models (Bhat et al. 2013; Osswald et al. 2015; Venkatesh et al. 2015, 2017; Venkataramani et al. 2019) and many previous clinical trials (Tan et al. 2020), no effective therapies exist for patients (Le Rhun et al. 2019). Unfortunately, patients diagnosed with GBM survive on average for only 12-24 months after the initial diagnosis (Tykocki and Eltayeb 2018). It is believed that inter- and intra-tumoral heterogeneity (Patel et al. 2014; Hu et al. 2017; Skaga et al. 2019; Wenger et al. 2019) as well as the brain microenvironment (Venkatesh et al. 2015, 2019; Monje et al. 2020) may be the factors influencing clinical trial outcomes.

GBM heterogeneity has been characterized according to histopathological tumor appearance, marker genes, and single-cell RNA sequencing (scRNA-seq) (Huse et al. 2011; Neftel et al. 2019; Bhaduri, di Lullo, et al. 2020; Couturier et al. 2020a). The GBM molecular subtypes based on genomic and transcriptional profiling include: pro-neural (PN), neural (N), classical (C), and mesenchymal (MES) (Balasubramaniyan et al., 2015; Bhat et al., 2013; Phillips et al., 2006). However, multiple molecular subtypes can coexist within a tumor at different relative preponderance and change identity over time (Sottoriva et al. 2013; Patel et al. 2014; Wang et al. 2018). The scRNA-seq profiling of GBM cells has allowed tumor cell classification based on the expression of cell identity markers into mesenchymal (MES)-, oligodendrocyte progenitor (OPC)-, astrocyte (AC)-, and neural progenitor cell (NPC)-like tumor cells (Neftel et al. 2019). Specific genetic aberrations primarily drove this classification, but the knowledge of genetic aberrations remains insufficient to predict tumor responses to specific treatments (Jacob et al. 2020).

The effects of the brain microenvironment on GBM cells have recently been investigated using human stem cell-derived brain organoids GBM (Bian et al. 2018a; da Silva et al. 2018a; Ogawa et al. 2018a; Linkous et al. 2019a; Pine et al. 2020a; Tang et al. 2020). Organoids recapitulate many important aspects of early human brain development (Eiraku et al. 2008; Mariani et al. 2012; Kadoshimaa et al. 2013; Lancaster et al. 2013; Pasca et al. 2015; Qian et al. 2016), including organization and the presence of primate-specific neural stem cells, [i.e., outer radial glia (oRG)] (Kadoshimaa et al. 2013; Qian et al. 2016; Watanabe et al. 2017), that have been associated with the expansion of the human cerebral cortex (Rakic 2009; Lui et al. 2011) and disorders such as lissencephaly (Bershteyn et al. 2017) and GBM (Bhaduri, di Lullo, et al. 2020). Brain organoids and GBM cells grown in suspension in defined media (GBM spheroids) can efficiently assemble into organoid-GBM chimeras. Tumor cells can invade organoids and proliferate without requiring exogenous growth factors (Lee et al. 2006a). Importantly, it has also been demonstrated that GBM cells in chimeras possessed increased resistance to irradiation and chemotherapy (Linkous et al. 2019a) and maintained single-cell gene expression signatures reminiscent of those detected in the resected primary parental tumors (Pine et al. 2020a). However, the molecular signaling pathways activated in GBM cells by fusion with organoids remain unknown. In addition, it is still unclear whether and how tumor cells influence the development of normal neural cells in organoids.

In this study, we demonstrated that mesenchymal and pro-neural GBM spheroids could be efficiently propagated without the tumor-required growth factors when fused with telencephalic organoids consisting of subpallial neural progenitors and GABAergic inhibitory neurons. We used single-cell RNA sequencing to investigate gene expression profiles in single cells obtained from GBM spheroids, organoids, and organoid-GBM chimeras to gain insights into the molecular pathways associated with tumor growth in the human brain-like environment. We found that both mesenchymal and pro-neural GBMs suppressed the gene expression network associated with telencephalic development and neuronal differentiation in organoids. On the other hand, organoids suppressed the expression levels of mitochondrial and proliferation genes and upregulated those associated with cell communication, neuronal activity, angiogenesis, and hypoxia in GBMs. Overall, this study describes genes and signaling pathways associated with tumor growth and propagation in the organoid-GBM chimeras. It also provides insights into the effects of GBM tumors on normal human brain development.

## RESULTS

### Multiple neural and mesenchymal cell types coexist in patient-derived GBM spheroids

Patient-derived GBM spheroids have been used as *in vitro* models for studying the cellular and molecular changes in human brain tumors and for drug discovery (Lee et al. 2006b; Ledur et al. 2017). However, the propagation and survival of GBM tumors *in vitro* are often dependent on the presence of exogenous epithelial growth factor and basic fibroblast growth factor (EGF and FGF, respectively) in the culture media (Lee et al. 2006b). EGF and FGF are well-known morphogens that can influence cell identity, gene expression profile, and resistance to anti-cancer treatments (Szymczyk et al. 2021). We investigated the properties of mesenchymal (MS) and proneural (PN) GBM spheroids obtained from patient-derived brain tumors using scRNA-seq (**Fig. 1A-B**) (Bhat et al. 2013; Balasubramaniyan et al. 2015). Neither MS- or PN-GBM spheroids survived for more than 12 days without EGF/FGF *in vitro* (**Fig. 1C-F**), which is consistent with the results of the previous study on these tumors (Bhat et al. 2013). However, both PN- (GBM11) and MS (GBM20) spheroids were robustly propagated in the presence of EGF and FGF (**Fig. 1C-F**).

**Figure 1.**
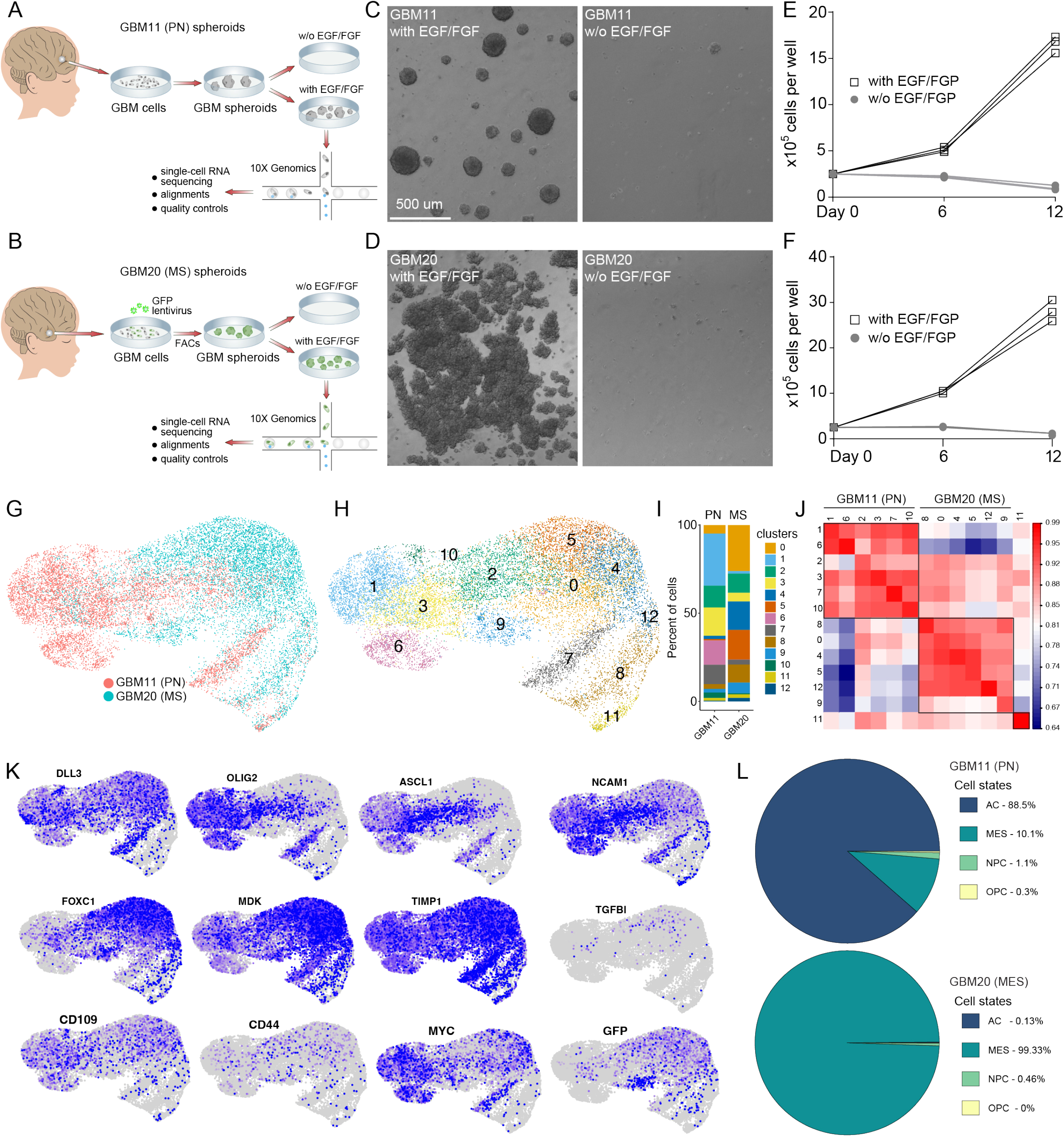
**A-B** Schematics of the pipeline for the derivation, different culturing conditions and single-cell profiling of GBM11 (PN) and GBM20 (MS) GBMs. **C-D** Images of GBM11 (C) and GBM20 (D) spheroids grown in presence (left) or absence (right) of exogenous EGF-FGF supplementation. **E-F** Growth curves for GBM11 (E) and GBM20 (F) spheroids with or without EGF/FGF. **G** UMAP plot of clusters from GBM20 and GBM11 cells in spheroids. **H** UMAP plot of Seurat clusters. **I** Bar plot showing the cluster composition of PN and MS tumors in spheroids. Stacked bars show the proportional composition for each tumor. **J** Similarity matrix of clusters from GBM11 and GBM20 tumor cells. Heatmap color indicates the Spearman ranked correlation. **K** Feature plots for selected PN, MES and cancer stem cell genes in spheroids. **L** Pie charts of percentages of cells in spheroids classified as astrocytic (AC)-like, mesenchymal (MES)-like, neural-progenitor (NPC)-like, and oligodendrocyte-progenitor (OPC)-like states. Scale bars: 500 μm (C-D). Data are presented as mean ± standard error of the mean (s.e.m.).

To compare and contrast the cellular composition in PN- (GBM11) and MS-(GBM20) spheroids, we collected single cells from multiple spheroids propagated in the presence of exogenous EGF (20 ng/mL) and FGF (20 ng/mL) for 12-14 days *in vitro* (**Fig. 1A-B**). We found substantially different cell compositions in these spheroids based on non-supervised cell clustering and expression levels of cell cluster-specific genes (**Fig. 1G-I**, **Supplementary Fig. 1**, and **Supplementary Dataset 1**). PN-GBM spheroids (8,421 cells analyzed) were primarily distributed between 6 cell clusters (**Fig. 1J**) and consisted of cells expressing pro-neural cell markers, including *ASCL1*, *OLIG2*, and *NCAM1* (**Fig. 1K**). MS-GBM spheroids (8,658 cells analyzed) were enriched with cells expressing mesenchymal cell markers, including *FOXC1, MDK*, and *TGFBI* (**Fig. 1K**), and distributed to a different set of 6 cell clusters (**Fig. 1J**). Both types of spheroids demonstrated an elevated expression of GBM stem cell markers *CD109* and *MYC*, PN-GBM marker *DLL3*, and MS-GBM markers *TIMP1* and *CD44* (Bhat et al. 2013)(**Fig. 1K**). Interestingly, tumor cells in both MS- and PN-GBM spheroids demonstrated increased expression of genes associated with subpallial telencephalic cell identities, including *FOXG1*, *SOX3*, *DLX1/2*, and *ASCL1* (**Supplementary Fig. 2**), suggesting the subpallial developmental origin of these tumors (Couturier et al. 2020b).

To further characterize the cell composition of MS- and PN-GBM spheroids, we assessed the identities of tumor spheroid cells using the average expression of gene signatures associated with different cellular states detected in primary GBMs, such as NPC-, AC-, OPC- and MES-like cell states (Neftel et al. 2019). This analysis revealed that PN-GBM spheroids primarily consisted of AC-like cells (88.6%) with smaller fractions of MES- (10.1%), NPC- (1.1%), and OPC-like cells (0.27%) (**Fig. 1L**). In contrast, MS-GBM spheroids were composed of MES-like cells (99.33%) with minor fractions of NPC- and AC- (0.46% and 0.13%, respectively) and no OPC-like cells (**Fig. 1L**). These findings highlight the distinct cellular composition and gene-expression profiles of PN-GBM and MS-GBM spheroids, underscoring their unique identities. However, despite these differences, the in vitro growth of both PN-GBM and MS-GBM spheroids depended on the addition of exogenous EGF and FGF to the culture medium. Since EGF and FGF are potent morphogens that regulate cell identities and proliferation, we next investigated whether GBM spheroids can be propagated without growth factors in co-culture with human stem cell-derived telencephalic organoids to mimic a human-brain-like microenvironment of GBM tumors in the developing brain.

### Generation and characterization of hPSC-derived subpallial telencephalic organoids from isolated single neural rosettes

We previously demonstrated that telencephalic organoids generated from isolated human stem cell-derived single neural rosettes (SNR) consist of pallial and subpallial neural progenitors (NPs) and both excitatory and inhibitory neurons and show reproducible cellular composition and organization (Wang et al. 2022). To create a microenvironment appropriate for co-culturing GBMs that exhibit the expression of subpallial telencephalic markers, including FOXG1, NKX2.2, SOX6, and DLX1 (**Supplementary Fig. 2**), we adapted the protocol to generate subpallial telencephalic organoids from SNRs (**Fig. 2, Supplementary Fig. 3,** and **Supplementary Dataset 2)**. SNR-derived subpallial organoids were generated in three separate batches from two different human pluripotent stem cells (hPSC) lines: embryonic stem cell (ESC) line H9 and control induced pluripotent stem cell (iPSC) line (AICS-0054-09), in which membrane-tagged RFP was inserted into the AAVS1 locus under the control of the CAGGS promoter (RFP-iPSC). Consistent with our previous observations (Wang et al. 2022), we observed that SNR-derived organoids increased in size over time and maintained a single visible lumen (**Fig. 2A**). At about two weeks post-rosette isolation, SNR-derived subpallial organoids reached the size of 0.15 - 0.3 mm^2^ (**Fig. 2B**). To characterize the cellular composition of these organoids, we performed scRNA-seq and analysis on 17,893 cells collected from 12 organoids produced in 3 separate differentiation batches from H9 and iPSC-RFP lines, 3-9 organoids (2,190-15,703 cells) per batch (**Fig. 2C**). The cellular compositions of dissociated organoids were reproducible across differentiation batches (**Fig. 2C-E**). The majority of cells within the SNR-derived organoids exhibited expression of the telencephalic marker FOXG1 (**Fig. 2G-H**). The cells in clusters 3 and 11 displayed the expression of early subpallial neural progenitor (NP) markers, including *NES, SOX2, NKX2-1, GSX2*, and *NKX2-1*, indicating their identity as subpallial telencephalic NPs (**Fig. 2F-H**). Conversely, clusters 7 and 9 exhibited heightened expression levels of intermediate subpallial NP markers such as *ASCL1, DLX1/2*, and *LHX6*, identifying them as subpallial telencephalic intermediate progenitors (IPs) (**Fig. 2F-H**). The cells in the remaining clusters (0-2, 4-6, and 8) were characterized by the expression of markers associated with GABAergic neurons, including *STMN2, GAD1*, and *GAD2*, supporting their classification as inhibitory neurons (**Fig. 2F-H**). Importantly, we detected no or very low expression of the pallial telencephalic NP and IP, and glutamatergic cortical neuron markers, including *PAX6, EOMES, SLC17A7* and *TBR1* (**Fig. 2G-H** and **Supplementary Fig. 3**), which is consistent with the subpallial telencephalic identities of cells in these organoids. These results suggest that subpallial telencephalic organoids can be reproducibly generated from SNRs.

**Figure 2.**
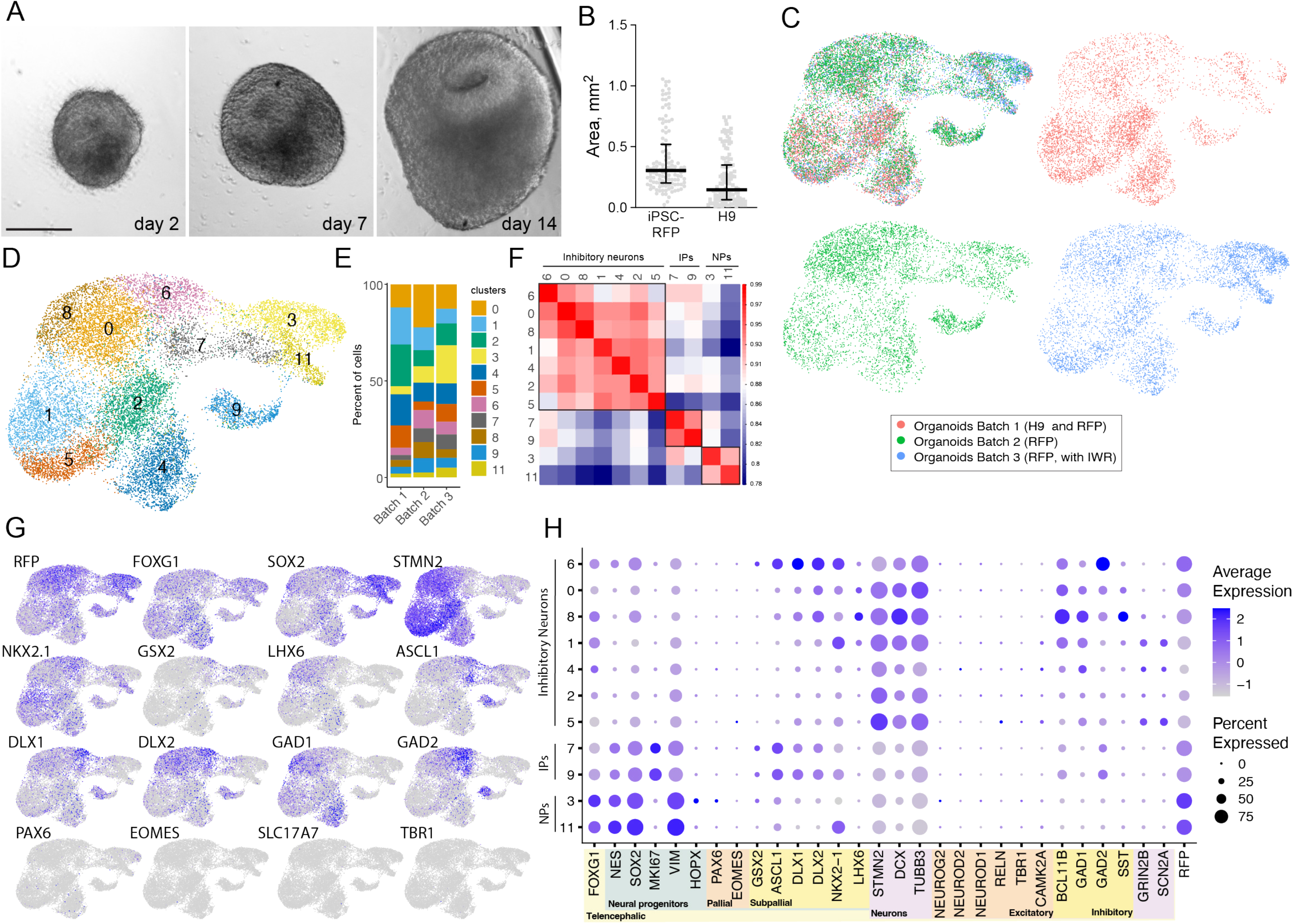
**A** Images of SNR-derived subpallial organoids at 2, 7, and 14 days in vitro post-isolation. **B** Scatter plot sized of organoids produced from iPSC-RFP and H9 stem cell lines in 3 (2 iPSC, combined and 1 H9) independent differentiation batches. Each dot represents an organoid. **C** UMAP plots of cells collected from multiple organoids obtained in 3 different differentiation batches from iPSC-RFP and H9 stem cell lines. **D** UMAP plot of Seurat clusters. **E** Bar plot showing cell cluster compositions for 3 different differentiation batches. **F** Similarity gene expression matrix for different cell clusters (IPs: intermediate progenitors; NPs: neural progenitors). **G** Feature heat-map plots for selected pallial and subpallial cell type-specific markers. **H** Dot plot for pallial and subpallial cell type-specific markers. Scale bar: 250 μm (A).

### Subpallial telencephalic organoids support *in vitro* propagation of GBM spheroids without exogenous growth factors

We assembled organoid-GBM chimeras by placing single MS- or PN-GBM spheroids together in the same well with a 1-month-old subpallial SNR-derived organoid (**Fig. 3A**). Both MS-GBM and PN-GBM spheroids exhibited 100% fusion rate with the organoids, resulting in the formation of organoid-GBM chimeras that demonstrated a progressive increase in size over time (**Fig. 3B** **and Supplementary Fig. 4**). This is consistent with the results of the previous studies on cerebral and cortical organoids and GBM spheroids (Bian et al. 2018b; da Silva et al. 2018b; Ogawa et al. 2018b; Linkous et al. 2019b; Pine et al. 2020b; Tang et al. 2020). Since cells in MS-GBM spheroids ubiquitously expressed GFP and SNR-derived organoids produced from the iPSC-RFP line expressed membrane-tagged RFP, we used this combination to investigate the relative distributions of tumor (GFP-expressing cells) and normal neural cells (RFP-expressing cells) in chimeras (**Fig. 3C-D** and **Supplementary Movie**). We observed that both tumor cells and normal cells exhibited efficient infiltration into the organoid and tumor tissue, respectively. These results suggest that human subpallial telencephalic organoids provide essential factors that sustain the survival and propagation of GBMs *in vitro*, without the reliance on exogenous EGF and FGF.

**Figure 3.**
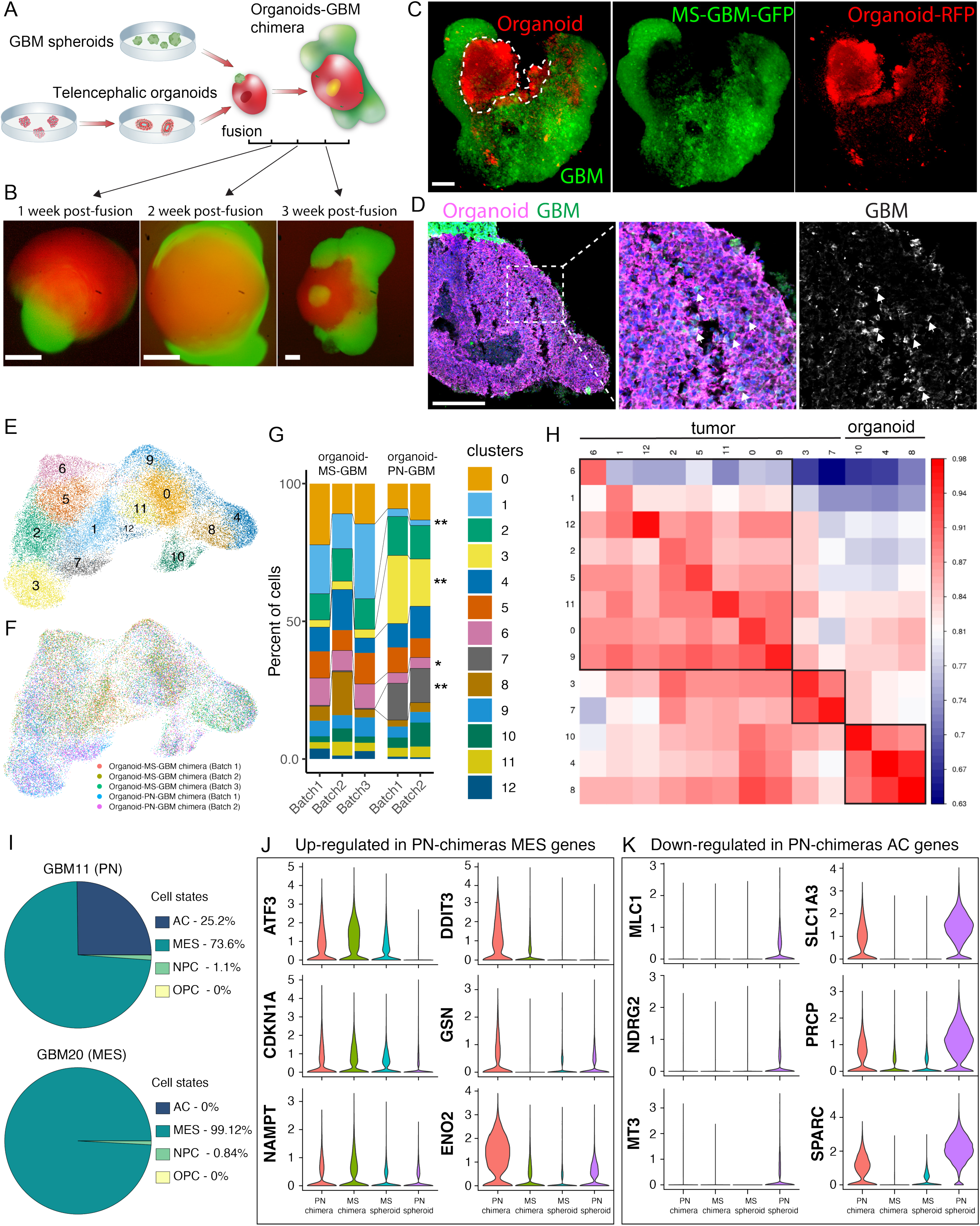
**A** Schematics of the experimental design for the generation of fused GBM spheroids and subpallial organoids (organoids-GBM chimeras). **B** Images of GBM20-iPSC chimeras at weeks 1, 2, and 3 post fusion. **C** Images of cleared MS-GBM-iPSC chimera. **D** Images of MS-GBM-iPSC chimera. **E** UMAP plot of Seurat clusters. **F** UMAP plot of cells coming from 5 independent groups of chimeras from 3 distinct batches. **G** Bar plot showing percentages of cell clusters in MS- and PN-chimeras. **H** Similarity matrix of clusters from chimeras. **I** Pie charts of percentages of tumor cells in MS- and PN-chimeras classified as AC-like, MES-like, NPC-like, and OPC-like states. **J** Violin plots of gene expression for selected MS and PN genes in tumor cells from chimeras and spheroids. Scale bar: 200 μm (B-D). Benjamini and Hochberg FDR: *<0.05, **<0.01.

To gain insights into the cellular composition of organoid-GBM chimeras and the molecular pathways altered in tumor cells by organoids, we performed single-cell RNA sequencing and non-supervised clustering analysis on 39,554 cells collected from 12 MS- and 6 PN-GBM-organoid chimeras that we generated in five separate differentiation batches (**Fig. 3E-G** and **Supplementary Dataset 3**). The analysis revealed similar compositions of cells in MS-GBM-organoid chimeras across 3 differentiation batches and in PN-GBM-organoid chimeras across 2 differentiation batches (**Fig. 3G**). However, the composition of cells in MS-GBM-organoid chimeras remained significantly different as compared to PN-GBM-organoid chimeras (**Fig. 3G**). In addition, analysis of gene expression profiles in the chimeras revealed distinct differences between GBM and organoid cells; therefore, these cells were largely distributed to different cell clusters (**Fig. 3H**). Notably, the distinction between PN-GBM and MS-GBM cells based on their gene expression profiles (tumor clusters) (**Fig. 3H**) was less apparent within chimeras compared to the separation observed in spheroids (**Fig. 1J**), suggesting that the organoid microenvironment likely influences the gene expression profiles of the tumor cells.

Consistent with this idea, we found that the composition of tumor cells in PN-GBM chimeras underwent significant changes, with a notable shift towards "MES-like" cell states (AC: 25.2%, MES: 73.6%, NPC: 1.1%, and OPC: 0% of all cells) (**Fig. 3I**), in contrast to PN-GBM spheroids (**Fig. 1L**). However, the cell composition in MS-GBM chimeras (MES: 99.12% and NPC: 0.84% of all cells) (**Fig. 3I**) remained similar to that observed in MS-GBM spheroids (**Fig. 1L**). This shift in composition was accompanied by the up-regulated expression of several MES-signature genes (Neftel et al., 2019), including *ATF3, CDKN1A, NAMPT, DDIT3, GSN,* and *ENO2* (**Fig. 3J**), and down-regulation of AC-signature genes, such as *MLC1, NDRG2, MT3, SLC1A3, PRCP,* and *SPARC* (**Fig. 3K**), in PN-GBM chimeras. This result supports the notion that tumor cell identities are plastic and can be significantly influenced by the surrounding microenvironment (Lee et al. 2008; Berezovsky et al. 2014; Neftel et al. 2019; Jung et al. 2021; Lauko et al. 2022; Yabo et al. 2022). Moreover, it implies that human brain-like conditions provided by subpallial telencephalic organoids facilitate the adoption of the MES-like gene expression signature by PN-GBMs. The MES-like GBM state is known to be associated with therapy resistance and poor prognosis for survival (Colman et al. 2010; Bhat et al. 2013).

### Tumor cells in organoids-GBM-chimeras undergo transcriptional changes consistent with altered glycolysis, quiescence, invasiveness, and angiogenesis

To understand the molecular changes occurring in tumor cells within chimeras, we compared the gene expression profiles of tumor cells from chimeras and those from spheroids (**Fig. 4A**). Non-supervised clustering analysis was performed on all cells collected from GBM spheroids, consisting of 8,658 cells from MS-GBM spheroids and 8,421 cells from PN-GBM spheroids, as well as non-RFP-expressing tumor cells collected from chimeras containing RFP-expressing organoids, including 7,797 cells from MS-GBM chimeras and 4,528 cells from PN-GBM chimeras (**Fig. 4B-C** and **Supplementary Dataset 4**). This analysis revealed that there were still substantial differences in the gene expression profiles between PN and MS tumor cells from both chimeras and spheroids. Indeed, PN tumor cells from chimeras exhibited elevated expression of pro-neural genes, including *OLIG2, ASCL1*, and *NCAM1* (**Supplementary Fig. 5**). In contrast, MS-GBM cells from chimeras showed increased expression of genes associated with mesenchymal cell states, such as *FOXC1, MDK*, and *FGFB1* (**Supplementary Fig. 5**). Consequently, PN tumor cells and MS tumor cells from both spheroids and chimeras were largely distributed to distinct cell clusters (**Fig. 4D-G**). However, the composition of cell clusters and the proportion of cells in these clusters were similar between PN -GBMs or MS-GBMs from chimeras and PN-GBMs or MS-GBMs from spheroids, respectively (**Fig. 4G**). Thus, PN tumor cells from spheroids were pooled with PN tumor cells from chimeras, and MS tumor cells from spheroids were pooled with MS tumor cells from chimeras for the identification of differentially expressed genes and molecular programs (**Fig. 4** **H-I** and **Supplementary Datasets 5-6**). This analysis revealed that several mitochondrial genes, including those encoding oxidative phosphorylation system (OXPHOS) complexes (*MT-ATP6*, *MT-C03*, *MT-CYB, MT-ND1, and MT-ND3)* and mitochondrial isomerase *EC1,* were significantly downregulated in both PN and MS tumor cells from chimeras as compared to those from spheroids (**Fig. 4H-J**). These changes are consistent with the idea that GBM cells in chimeras more readily utilize glycolysis for energy metabolism outside the mitochondria than GBM cells in spheroids. Indeed, several genes associated with glycolysis, including *DDIT3*, *PGK1*, and *NDRG1,* and mitophagy, including *BNIP3* and *BNIP3L,* were upregulated in GBM cells from chimeras (**Fig. 4K**). Other genes upregulated in chimeras were activity-regulated immediate early genes *EGR1*, *IER2*, and *FOS* (Sun and Lin 2016), angiogenesis gene *VEGFA* (Bao et al. 2006), tumor invasiveness and metastases gene *MALAT1* (Venkataramani et al. 2022), and hypoxia and oxidative stress genes *BNIP3L, FAM162A, BNIP3, HILPDA, PGK1, NDRG1, VEGFA,* and *DDIT3* (**Fig. 4K**). These results suggest that GBM cells exposed to a human brain-like environment adapt to it by altering the expression levels of multiple genes associated with cell metabolism. The observed changes may contribute to increased quiescence of tumor cells in chimeras based on increased glycolysis (Zhou et al. 2011; Strickland and Stoll 2017), invasiveness, and vascularization.

**Figure 4.**
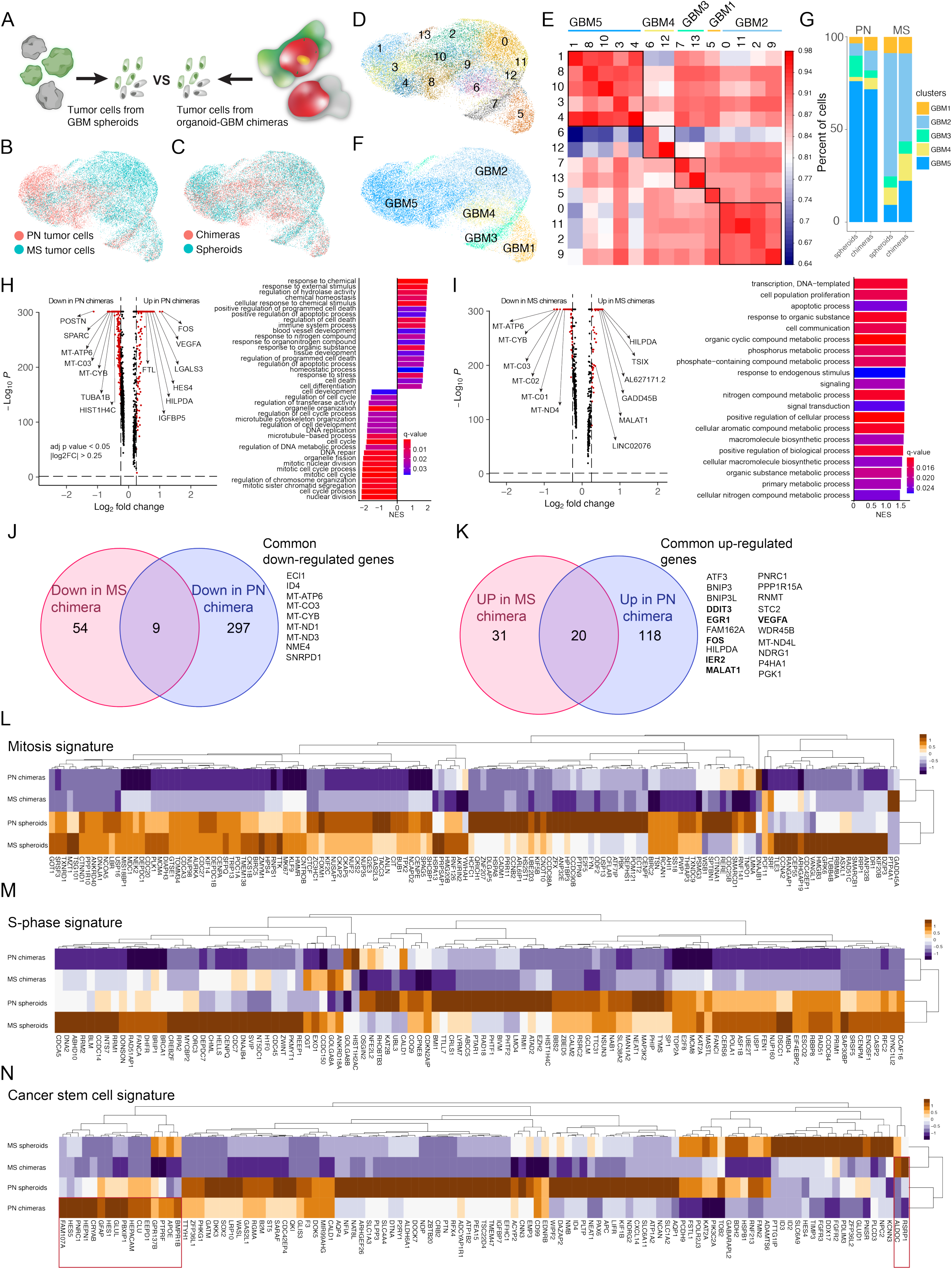
**A** Scheme of the differential expression analysis workflow. **B-C** UMAP plot of cells coming from tumors clustered by origin (chimeras or spheroids [B]) or tumor type (C). **D** UMAP plot of clusters from combined tumor cells coming from Chimeras and spheroids, colored by cluster. **E** Similarity matrix of clusters from combined analysis. Heatmap color indicates the Spearman ranked correlation, and above the heatmap is the annotated “grouped cluster” for each cluster. **F** UMAP plot of “grouped clusters”. **G** Bar plot showing the cluster composition of PN and MS tumors in Chimeras and spheroids. **H** Volcano plot of differentially expressed genes shows the most up- and down-regulated genes found in PN tumor cells in Chimeras versus spheroids (left). Gene sets significantly enriched (pvalue < 0.05) in the PN tumor grown in Chimeras using the Gene Ontology Biological Processes, ordered by NES [Normalized Enrichment Score] (right). A positive NES indicates enrichment in the PN tumor grown in Chimeras, a negative NES indicates enrichment in the PN tumor grown in spheroids. **I** Volcano plot of differentially expressed genes shows the most up- and down-regulated genes found in MS tumor cells in Chimeras versus spheroids (left). Gene sets significantly enriched (pvalue < 0.05) in the MS tumor grown in Chimeras using the Gene Ontology Biological Processes, ordered by NES (right). **J** Venn diagram showing the number of common (list on the right) and unique upregulated genes (adjusted pvalue < 0.05, logFC > 0.25) in PN and MS tumors in Chimeras. **K** Venn diagram showing the number of common (list on the right) and unique down-regulated genes (adjusted pvalue < 0.05, logFC < -0.25) in PN and MS tumors in Chimeras. **L-N** Heat maps of the “mitosis”, “s-phase”, and “cancer stem cell” gene signatures for PN and MS tumors grown in spheroids and chimeras.

To understand the effects of the organoid microenvironment on the proliferation/ quiescence of GBM cells in chimeras, we investigated the expression levels of marker genes associated with mitosis and cell cycle (Xie et al. 2022). We found dramatically reduced expression levels of the mitosis and S-Phase signature genes in both MS- and PN-GBM cells in the chimeras compared to spheroids (**Fig. 4L-M**). These results suggest that GBM cells in chimeras are less proliferative and possibly more quiescent than those in spheroids. However, we found that only certain glioblastoma quiescence signature genes [“cancer stem cell signature” (Xie et al. 2022)] were upregulated in PN-GBM (16 out of 118) and MS-GBM (2 out of 118) cells in chimeras (**Fig. 4N**). Interestingly, among the genes upregulated in PN-GBMs, some were outer radial glia (oRG) marker genes, such as *CLU*, *HES1*, *GFAP*, *CRYAB*, and *FAM107A* (Pollen et al. 2015). Elevated expression of oRG genes was identified in a population of oRG-like cancer stem cells in primary glioblastoma tumors (Bhaduri, di Lullo, et al. 2020). Together, these results suggest that GBM cells exposed to a human brain-like environment adapt by altering their gene expression profiles towards increased glycolysis, reduced proliferation, and increased invasiveness and angiogenesis, all of which are hallmarks of GBM within their native state *in vivo*.

### Neural cells in organoids-GBM-chimeras undergo transcriptional changes consistent with altered oxidative stress, angiogenesis, and neurodifferentiation

Our understanding of the effects of tumor cells on normal brain development is limited due to the inaccessibility of normal human brain tissue. We reasoned that comparing cell compositions and gene expression profiles of organoids and organoids in chimeras could provide unique insights into the effects of tumor cells on normal human brain development. To compare and contrast the properties of neural cells in chimeras and organoids, we co-clustered RFP-expressing organoid cells from chimeras (MS-GBM chimeras: 1,188 cells and PN-GBM chimeras: 2,295 cells) with those from H9 and iPSC-derived RFP-expressing organoids (17,893 cells) (**Fig. 5A-B** and **Supplementary Dataset 7**). This analysis revealed that the compositions of neural cell clusters and the proportions of cells in different clusters remained relatively similar between organoids and chimeras (**Fig. 5C-E**). The identified cell clusters could be broadly divided into the clusters of NPs (clusters: 4, 9, 10, and 11) and GABAergic neurons (clusters: 0, 1, 2, 3, 5, 6, 8, and 12) (**Fig. 5F**). To identify differentially expressed genes and molecular pathways between NPs and GABAergic neurons in organoids and chimeras, we performed a ‘pseudo-bulks’ mRNA-seq analysis (Squair et al. 2021) on the combined clusters of NPs and neurons (**Fig. 5F**). Both NPs and neurons in chimeras were different from those in organoids; however, the dissimilarity between the neurons in chimeras was significantly greater than that of the NPs (**Fig. 5G-H**). To identify dysregulated neuronal pathways, we performed GSEA analysis on DEGs identified in neurons (**Fig. 5I** and **Supplementary Dataset 8**) and NPs (**Fig. 5J** and **Supplementary Dataset 9**). The molecular pathways “*response decreased oxygen levels”* and *“response to hypoxia”* were upregulated in both neurons and NPs from chimeras as compared to those from organoids (**Fig. 5I-J**). This finding suggests that GBMs induce the formation of a hypoxic pro-tumorigenic environment in organoids. The hypoxic microenvironment has been associated with aggressive and therapy-resistant tumors (Ye et al. 2022) and with the abnormal specification of human cortical neurons (Bhaduri, Andrews, et al. 2020). In addition, we observed increased enriched molecular pathways “*positive regulation of vasculature development” and “angiogenesis”* in neurons, and upregulation of pro-angiogenesis gene *VEGFA* in NPs (**Fig. 5I-J**), suggesting that GBMs promote the formation of a pro-angiogenic hypoxic microenvironment in neural tissue by altering the gene expression profiles of neural cells.

**Figure 5.**
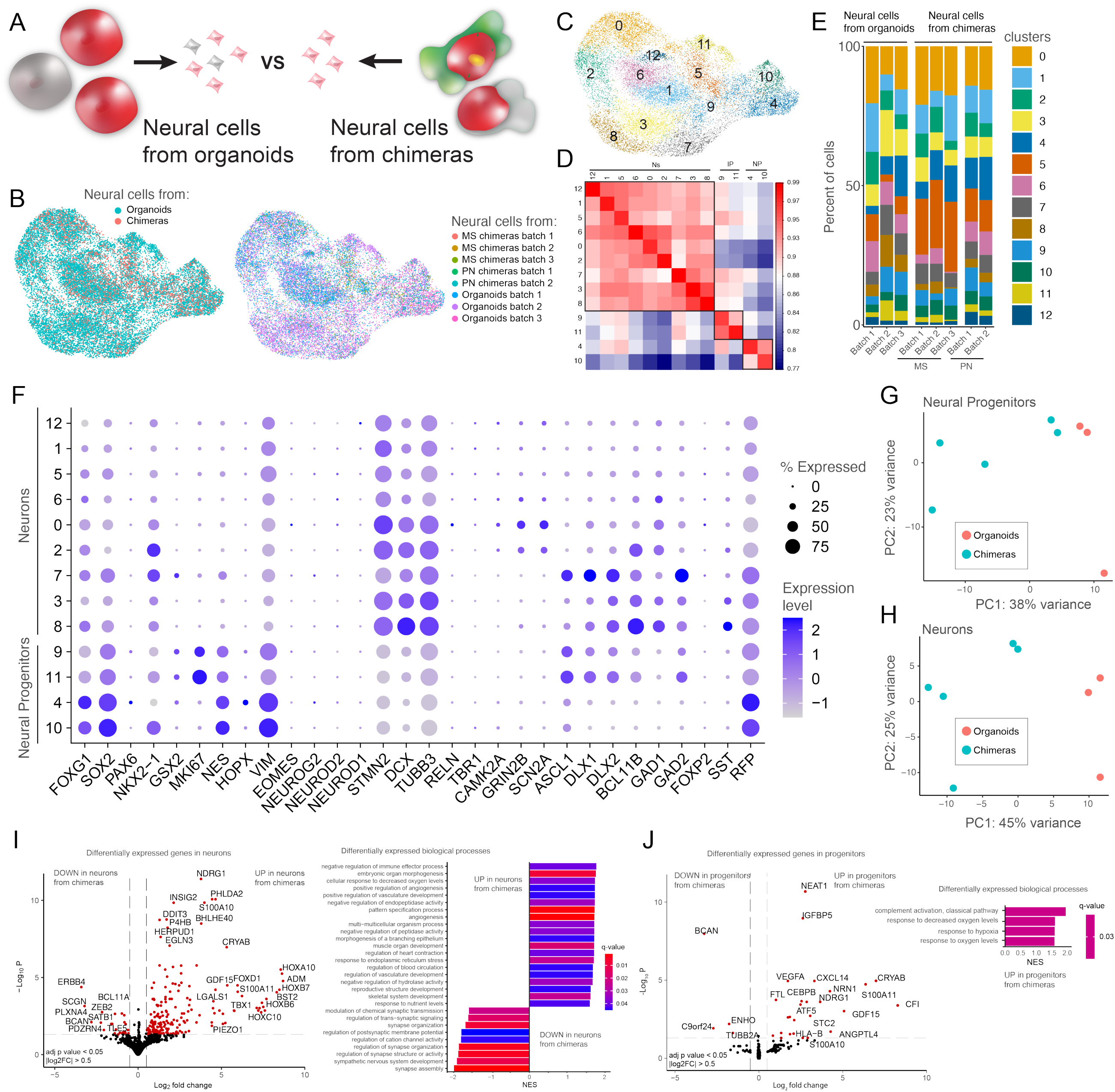
**A** Schematics for the differential expression analysis design. B UMAP plot of cells coming from organoids clustered by origin (organoids or chimeras, left) or sample name (right). C UMAP plot of Seurat clusters. D Similarity matrix of clusters from organoids and organoids in chimeras (MS and PN). E Bar plot showing the percentages of clusters in different batches of organoids and organoids in chimeras (MS and PN). F Dot plot for pallial and subpallial specific markers in organoids’ clusters. **G-H** Principal Component Analysis (PCA) plots of pseudo bulk RNA seq values for Neural Progenitors (G) and Neurons (H) in organoids and chimeras. **I** Volcano plot of differentially expressed genes shows the most up- and down-regulated genes found in Neuronal cells in Chimeras versus Organoids (left). Gene sets significantly enriched (pvalue < 0.05) in Neurons in Chimeras using the Gene Ontology Biological Processes, ordered by NES (right). **J** Volcano plot of differentially expressed genes shows the most up- and down-regulated genes found in Neural Progenitor cells in Chimeras versus Organoids (left). Gene sets significantly enriched (pvalue < 0.05) in Neural Progenitors in Chimeras using the Gene Ontology Biological Processes, ordered by NES (right).

In neurons from chimeras, the down-regulated molecular pathways were “*synapse assembly”*, “*synapse organization”,* and “*ion channel activity”* (**Fig. 5I**), suggesting potential impairments of neuronal differentiation and maturation of neurons in chimeras by tumor cells. This is consistent with a previous study that reported decreased inhibitory synapses in a patient-derived xenograft mouse model (Yu et al. 2020). Together, these results show that normal neural development is impacted by GBM tumors, which cause impaired neuronal differentiation and maturation and increased formation of a pro-tumorigenic environment, similar to what was previously observed in patients and xenotransplantation models (Osswald et al. 2015; Venkataramani et al. 2019, 2022; Hausmann et al. 2022).

## DISCUSSION

In this study, we fused patient-derived PN- and MS-GBMs with human subpallial telencephalic organoids to investigate the effects of the human brain-like environment on tumor cells and the effects of GBM tumors on human brain development. We observed efficient propagation of GBM tumors in chimeras without the need for exogenous growth factors and substantial changes in gene expression profiles in both tumor and neural cells. Collectively, our findings suggest that tumor and normal neural cells extensively interact with each other in organoid-GBM chimeras and that the use of organoid-GBM chimeras may provide a physiologically relevant microenvironment for propagating GBM cells in vitro for future mechanistic insights and therapy development.

The fusion of GBMs with human stem-derived organoids has been investigated in several previous studies (Bian et al. 2018b; da Silva et al. 2018b; Ogawa et al. 2018b; Linkous et al. 2019b; Pine et al. 2020b). These studies demonstrated that GBM cells can spontaneously penetrate the organoids and form interconnected networks of tumor and “normal” cells. Similar to the previous studies, we also observed an efficient fusion of GBMs with human stem cell-derived organoids. In contrast to those studies, we fused GBMs with well-defined (in terms of cellular composition) subpallial telencephalic organoids. Both PN- and MS-GBMs demonstrated an elevated expression of the subpallial telencephalic marker genes, suggesting that subpallial NPCs may give rise to GBM tumors (Sanai et al. 2005; Eichmüller et al. 2022) and provide a more supportive microenvironment for growing GBM tumors *in vitro*.

Unlike in GBM spheroids, we observed that tumor cells in organoid-GBM chimeras efficiently propagated without supplementation of exogenous EGF and FGF. EGF and FGF are commonly used for propagating GBM spheroids *in vitro* (Lee et al. 2006b). These factors promote tumor cell survival and proliferation by activating the MAPK/ERK and PI3K/AKT/mTOR signaling pathways (Schlessinger 2000). However, the physiological concentration of these factors in the brain is largely unknown and may vary depending on the brain region, microenvironmental niche, age, physiological state, or pathological state. We found that GBM tumors showed limited propagation in spheroids without exogenous EGF and FGF but robust propagation in chimeras. These results suggest that either normal neural cells in organoids provide GBMs with EGF and FGF or alternative signaling cascades are activated in tumor cells for survival and propagation without these growth factors. As we also found no The specific signaling cascades activated in tumor cells by organoids remain to be understood.

Differentially expressed genes detected in GBM tumors in chimeras compared to those in spheroids (**Fig. 4**) may provide insights into the molecular programs involved in promoting tumor cell survival and propagation in chimeras. Among the upregulated genes in chimeras, we found transcription factor *ATF3*, associated with stress responses and more aggressive tumor behavior with poor prognosis (Xu et al. 2022); mitophagy regulators *BNIP3* and *BNIP3L* (Ney 2015), activated with hypoxia (Jung et al. 2019), which is a common feature of the tumor microenvironment (Zhou et al. 2011); and the MAPK/ERK pathway genes *FOS*, *DDIT3*, and *VEGFA* (Bao et al. 2006; Venkatesh et al. 2015), which are known to promote tumor cells survival and propagation. This suggests that oxidative stress induced by hypoxia and the MAPK/ERK pathway may be involved in supporting tumor cells in organoid-GBM chimeras.

Interestingly, we also found that GBM tumors in chimeras showed dramatically reduced expression of multiple genes associated with cell cycle proliferation (**Fig. 4**). Reduced proliferation is a hallmark of quiescent stem cells (Hanahan 2022). The quiescent tumor stem cells are characterized by slow rates of division (Antonica et al. 2022) and, thus, are difficult to target with irradiation or chemotherapies (Xie et al. 2022). Interestingly, it was demonstrated that GBM cells in organoid-GBM chimeras are substantially more resistant to irradiation (Linkous et al. 2019b). Our results suggest that the reduced proliferation of GBM tumors in chimeras is potentially responsible for this radioresistance. Interestingly, we found no increase in the expression levels of multiple quiescence genes in chimeras. Instead, different sets of quiescence genes were upregulated in different GBM tumors under different experimental conditions (**Fig. 4N**). These results suggest that the tumor-specific microenvironment may be involved in defining the tumor-specific quiescence signature and that alternative culture conditions such as organoid-GBM chimeras could be useful for studying the pathology of GBMs in the brain.

Another interesting observation of this study was that the co-culture of PN-GBM with subpallial telencephalic organoids that consist of GABAergic NPCs and neurons induced gene expression changes in PM-GBM consistent with the acquisition of mesenchymal cell identity (Bhat et al. 2013). PN to MS transition (PMT) in GBM has been previously observed in response to NF-kB exposure (Bhat et al. 2013), serum exposure (Balasubramaniyan et al. 2015), TGFβ exposure (Joseph et al. 2014), activation of the Notch signaling pathway (Cheng et al. 2017), and mutations in tumor suppressor genes such as PTEN and P53 (Behnan et al. 2019). In addition, changes in the tumor microenvironment, including hypoxia and inflammation, have been reported to promote the PMT microenvironment (Tejero et al. 2019). PMT is associated with increased tumor cell motility, invasiveness, and therapy resistance (Bhat et al. 2013). Our results support the idea that the environment in organoid-GBM chimeras promotes PMT. However, additional functional experiments are required to confirm the mesenchymal-like identities of PN-GBM tumors in chimeras.

Finally, in this study, we also investigated the effects of tumor cells on gene expression profiles of neural cells in organoids. The detected gene expression changes are consistent with impaired neural development and functional maturation, as well as increased angiogenesis and response to hypoxia (**Fig. 5**). Interestingly, a recent study showed that cell stress impairs the specification of oRG cells and neurons in organoids (Bhaduri, Andrews, et al. 2020), suggesting that the hypoxic stress affects the normal development in organoid-GBM chimeras. Moreover, the effect of GBMs on GABAergic neurons in organoid-GBM chimeras may lead to impaired function and elevated excitability due to the disinhibition of excitatory neurons and neural networks. Indeed, elevated excitability was observed in a recent study where authors demonstrated that GBM cells express high levels of IGSF3, an interactor of the potassium channel Kir4.1 that suppresses potassium buffering leading to spreading depolarization (Curry et al. 2023). Finally, another recent study showed increased hyperexcitability, increased PSD95^+^ excitatory synapses, and decreased Gephyrin^+^ inhibitory synapses in the peritumoral regions of mice transplanted with patient-derived GBM cells (Yu et al. 2020). Together, these results support the idea that GBM cells have a profound effect in remodeling the peritumoral neuronal network towards a state of hyperexcitability, with a selective detrimental effect on inhibitory synapses.

In summary, we demonstrated an efficient propagation of GBM tumors *in vitro* by fusing them with subpallial telencephalic organoids in chimeras. We also found that GBM cells adapt to the developing human brain microenvironment by altering gene expressions associated with the PN to MS transition, proliferation, hypoxia, and angiogenesis. In addition, the presence of GBM cells impacted the properties of neural cells in the organoids that were consistent with impaired neuronal differentiation and increased angiogenesis. Collectively, this study provides important insights into the molecular pathways that could be targeted for therapy development. Future work will be needed to explore cell-cell communications between organoids and GBM tumor cells to elucidate the underlying cellular and molecular mechanisms. Finally, future studies should also investigate the contributions of different types of NPCs, inhibitory and excitatory neurons, and different types of glial cells on GBM tumor survival, propagation, and treatment resistance.

## MATERIALS AND METHODS

### Cell lines

The GBM cell lines used in this study are: GBM11 (PN), GBM established cell line (Bhat et al., 2013) and GBM20 (MS), GBM established cell line (Bhat *et al*., 2013). The hPSC lines used in this study are: mTagRFP-T hiPSCs (Allen Cell Collection, Cat # AICS-0054-091) and H9 hESCs (WiCell). Both hPSC lines were obtained from healthy donors.

### Generating Single Neural Rosette-derived Organoids

The iPSCs were grown in the StemFlex/E8 medium (1:1 mixture of StemFlex (ThermoFisher, Cat # A3349401) and E8 (ThermoFisher, Cat # A3349401) media). The iPSCs were differentiated in the N2/B27 medium containing a 1:1 mixture of DMEM/F12 and Neurobasal-A, 1X N2 supplement (ThermoFisher Scientific), 1X B27 supplement with vitamin A (ThermoFisher Scientific), 2 μg/ml of Heparin, 1X GlutaMAX (ThermoFisher Scientific), and 1X MEM-NEAA. The N2/B27 medium was supplemented with 4 µM Dorsomorphin (Tocris, Cat # 3039) and 10 µM SB431542 (Tocris, Cat # 1614). SNR-derived organoids were generated as previously described (Wang et al., 2022). Briefly, the differentiated iPSCs were grown to rosettes in N2/B27 medium for 6 days. Single neural rosettes were picked and grown to single neural rosette-derived organoids in N2/B27 medium with 10 ng/ml of each EGF and FGF for 1-3 days without agitation and then with agitation for 2-3 weeks.

### Growing GBM Spheres

GBM11 and GBM20 spheres were growing as described previously (Bhat *et al*., 2013). Briefly, GBM cells were incubated in a nontreated tissue culture flask in the GBM spheres medium containing DMEM/F12 Medium, 2X B-27 supplement (Life Technologies, Cat # 17504-044), 1X Normocin (InvivoGen, Cat # ant-nr-1), 20 ng/ml EGF (Life Technologies, Cat # PHG0311) and 20 ng/ml FGF (Life Technologies, Cat # 13256-029). To maintain the proliferation, spheres must be triturated to single cells/small clumps once visible spheres appear.

### Generating Organoid-GBM Chimeras

To generate organoid-GBM chimeras, 2-3 week-old SNR-organoids were individually transferred to a V-shaped non-treated 96-well plate containing N2/B27 medium (no EGF/FGF), one organoid per well. 2-4 GBM spheres were added to each organoid/well. Organoid-GBM chimeras were incubated in a 37°C/5% CO_2_ incubator for 12-14 days with media change every other day. Organoid-GBM chimeras were then transferred to 24-well non-treated plates and maintained in a 37°C/5% CO_2_ incubator with agitation (∼45rpm) on an orbital shaker for one week. The media were changed every other day.

### Immunohistochemistry and confocal imaging

1 month old single neural rosettes (SNR)-derived organoids were fixed in 4% paraformaldehyde overnight at 4°C and washed with PBS and the organoids were then passed through a sequential sucrose solution gradient of 10%, 20% and 30% respectively. Organoids were placed in cryosectioning molds, embedded in OCT and flash frozen and stored at -80°C. Then organoids were cut at 15-20 μm thickness using a Leica Cryostat machine and adhered to positively charged microscope slides. Cryosections were then processed for immunostaining as follows: Sections were washed with 0.1M glycine in PBS and then blocked with 0.3% Triton-X 100 with 3% Bovine cerium albumin (BSA) in PBS for 1-2 h at room temperature. Primary antibodies were diluted in 0.3% Triton-X 100 with 3% BSA in PBS and left overnight at cold room. The following primary antibodies were used: mouse anti-PAX6 (DSHB, 1:50), sheep anti-TBR2 (R&D Systems, AF6166, 1:200), mouse anti-MASH1 (Invitrogen, 1:200), rat anti-CTIP2 (Abcam, ab18465, 1:200), rabbit anti-GSX2 (Invitrogen, PA5-35887, 1:200), mouse anti GAD67 (Millipore, MAB5406, 1: 200), mouse anti-SOX2 (Santa cruz, sc-365823, 1:200), rabbit anti-RFP (Evrogen, AB234, 1:1000), chicken anti-GFP (Novus, NB100-1614, 1:500). The next day, sections were washed with PBS for 15 min and secondary antibodies were also diluted with 0.3% Triton-X 100 with 3% BSA in PBS. The secondary antibodies coupled to Alexa Fluor fluorescent dyes (Invitrogen) were added 1:1000 ration in combination with DAPI (1:1000) for 1-2 h at room temperature in the dark. After 15 min PBS washes, slides were mounted with coverslips using Aqua Poly Mounting Media, left to dry at room temperature in the dark and stored at -20°C for imaging. Fluorescent imaging was performed using Zeiss 880 Airyscan confocal microscopes using 10x and 20x objective at the University of Utah Cell Imaging Core Facility and data processing and quantification was performed using Fiji, and/or CellProfiler software.

### Tissue clearing and confocal imaging

Organoids-GBM chimeras were fixed with fresh 4% paraformaldehyde (PFA) (Cat# 15710) for 24 hours at 4°C. The fixed organoids were then cleared following a modified SWITCH protocol developed by the Chung Lab (Murray et al. 2015). Briefly, chimeras were transferred to a fixation-OFF solution and incubated at 4°C for 1 day. Subsequently, chimeras were transferred to fixation-ON solution and incubated for 1 day at 4°C. To inactivate the fixative solution, chimeras were washed twice with PBST at room temperature for 1 hour each and then submerged into fixative-inactivation solution at 37°C overnight. Subsequently, chimeras were washed twice with thermal clearing solution for 1 hour each at 37°C. To achieve complete lipid clearing, chimeras were immersed in fresh thermal clearing solution and incubated overnight at 37°C. Finally, to achieve maximal optical clearing, chimeras were placed in 500 µL refracting index matching solution (RIMS) at 37°C for 1 hour. All incubations and wash volumes used were 10 mL, incubation conditions were stationary and in the dark to preserve chromophores. For imaging, chimeras were mounted on a glass slide using 1 mm thick homemade silicone ring spacer, filled with RIMS, and covered with a glass cover slip to avoid drying up and air bubbles. Fluorescent imaging was performed at the University of Utah Cell Imaging Core Facility using a Zeiss LSM 880 Airyscan confocal microscope equipped with a Plan-Neofluar 10X/0.30 M27 objective, 488 nm and 568 nm laser to acquire high-resolution images. We used a refractive index of 1.46 and a total of 64 Z-stacks were acquired to cover the entire thickness of the sample. To create 3D reconstructions and a video, we used Imaris Cell Imaging (Bitplane) and Fiji softwares.

### Single-cell collection from organoids and GBM-organoids chimeras

SNR-derived organoids were rinsed with sterile Dulbecco’s phosphate-buffered saline (DPBS, 1X) and chopped in small pieces using a scalpel. Organoids’ pieces were collected in 1.5 mL tubes in DPBS and centrifuged for 2 minutes at 1000 rpm. Samples were incubated in pre-warmed papain (Worthington Biochemical) containing 5% DNaseI (Sigma) at 37°C for 50 minutes. After incubation, cells were resuspended in Trypsin Inhibitor (Sigma) and manually triturated by pipetting up and down using P1000 pipette. Cells were centrifuged for 5 minutes at 1000 rpm before the next step. Cells were resuspended in cold DPBS containing 0.04% BSA. Samples were kept on ice from this point forward. Samples were quickly transferred to the High Throughput Genomics Core Facility at the University of Utah for cell viability assessment.

### Library preparation and raw data processing for single-cell RNA-seq

Single-cell capture, lysis, and cDNA synthesis were performed with the 10x Genomics Chromium at the High Throughput Genomics Core Facility at the University of Utah. Single cells were loaded onto 10x Genomics Single Cell 3’ Chips along with the master mix following the manufacturer’s instruction for the Chromium Single Cell 3′ Library (v3.1). The resulting libraries were sequenced on an Illumina NovaSeq 6000 in a 150 × 150 cycle paired-end sequencing mode using a NovaSeq 6000 S4 reagent Kit v1.5. Pre-processing of reads was conducted using the Cell Ranger pipeline (10x Genomics) with alignment to GRCh37 (hg19) reference assembly built with the addition of GFP and RFP sequences. Gene-barcoded matrices were constructed for each sample by counting unique molecular identifiers (UMIs). The single-cell RNA-seq data reported in this paper is available in the GEO repository with the following accession number (to be provided).

### Clustering analysis and removal of stressed cells in organoids

Downstream analysis of single-cell RNA-seq matrix files was conducted using Seurat (v 4.2.0 and R version 4.1.2). Cells were selected and filtered out based on QC metrics distributions across samples of the same experimental condition (number of unique genes detected in each cell [cells with less than 200 genes detected were filtered out], total number of molecules detected in each cell, and percentage of reads that map to the mitochondrial genome). Single cell expression values were normalized, and the top 2000 highly variable genes were identified using *FindVariableFeatures* function. Data were then scaled, and principal component analysis (PCA) was performed using 50 dimensions. Data integration between different samples was performed with Harmony algorithm. The *ElbowPlot* function was then used to pick the relevant dimensions. The top 10 principal components were used for running UMAP dimensional reduction technique and for clustering using Seurat’s *FindNeighbors* and *FindCluster* functions (original Louvain algorithm, resolution 1.0). Cluster identities were assigned based on (I) cluster gene markers as determined by *FindAllMarkers* function, (II) gene expression values of known telencephalic or GBM markers, and (III) cluster correlation as determined by *clustify* function (Spearman ranked correlation, clustifyr package v 1.6.0). Identification of stressed cells in organoids was performed using the package *gruffi* (v. 0.7.2) using default parameters and without adjusting manually the thresholds estimated by *gruffi*. Stressed cells were removed from the dataset before further analysis.

### Differential expression analysis

Differential gene expression analysis between clusters in different samples was performed using the *FindMarkers* function (Wilcoxon rank sum test) and genes with an adjusted (Bonferroni’s correction) p-value < 0.05 were considered significant. Volcano plots of significant differentially expressed genes were generated using the *EnhancedVolcano* function (|FoldChange| > 0.5 and adjusted pvalue < 0.05) from the EnhancedVolcano package (v. 1.12.0). Differential gene expression analysis between different samples was performed calculating pseudo bulk values using first the *SingleCellExperiment* function (SingleCellExperiment package, v. 1.16.0) to convert Seurat objects and then the *aggregate.Matrix* function (fun = “sum”, package Matrix.utils v. 0.9.8) to generate pseudo bulk RNA-seq datasets. Pseudo bulk datasets were passed to the *DESeq* function (DESeq2 package, v. 1.34.0) to calculate differentially expressed genes.

### Cell state scoring

Cell state scores were calculated using the lists of gene signatures (NPC-like, OPC-like, AC-like, and MES-like) previously reported in Neftel et al. 2019. For each tumor cell a signature score value was calculated using the *AddModuleScore_UCell* function from the UCell package (v.). The NPC, OPC, AC or MES identity was assigned to each cell by looking at the maximum signature score value. The pie charts with the percentages of each cell state per GBM type were generated using the *pie* function.

### Gene Set Enrichment Analysis

Gene Set Enrichment Analysis was performed using the *gseGO* function from the clusterProfiler package (v. 4.2.2). Gene-concept networks were generated using the *cnetplot* function from the enrichplot package (v. 1.14.2).

## Supplementary Figures

**Supplementary Figure 1.**
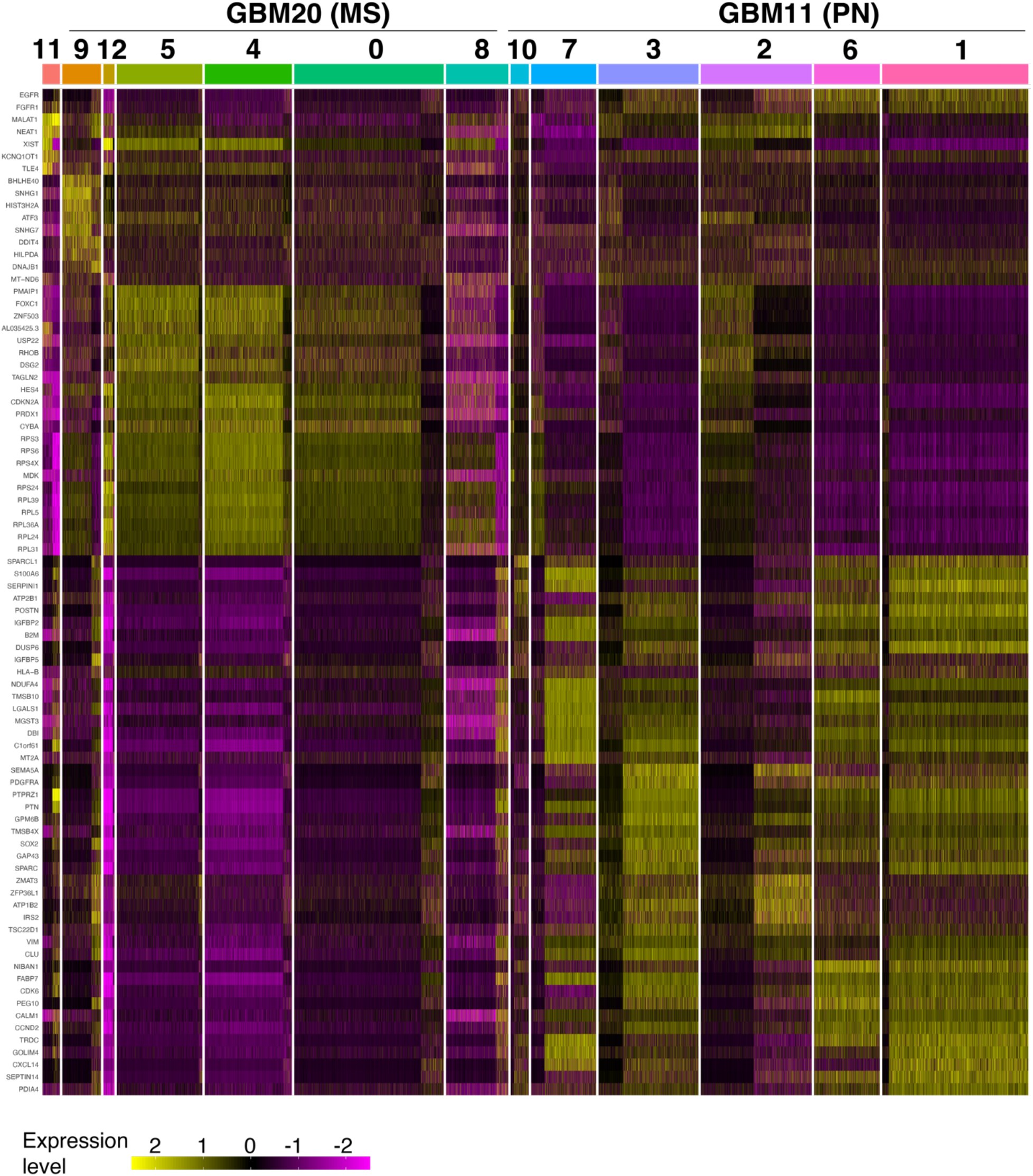
Heatmap of cluster-specific genes calculated in Seurat clusters from PN (GBM11) and MS (GBM20) spheroids.

**Supplementary Figure 2.**
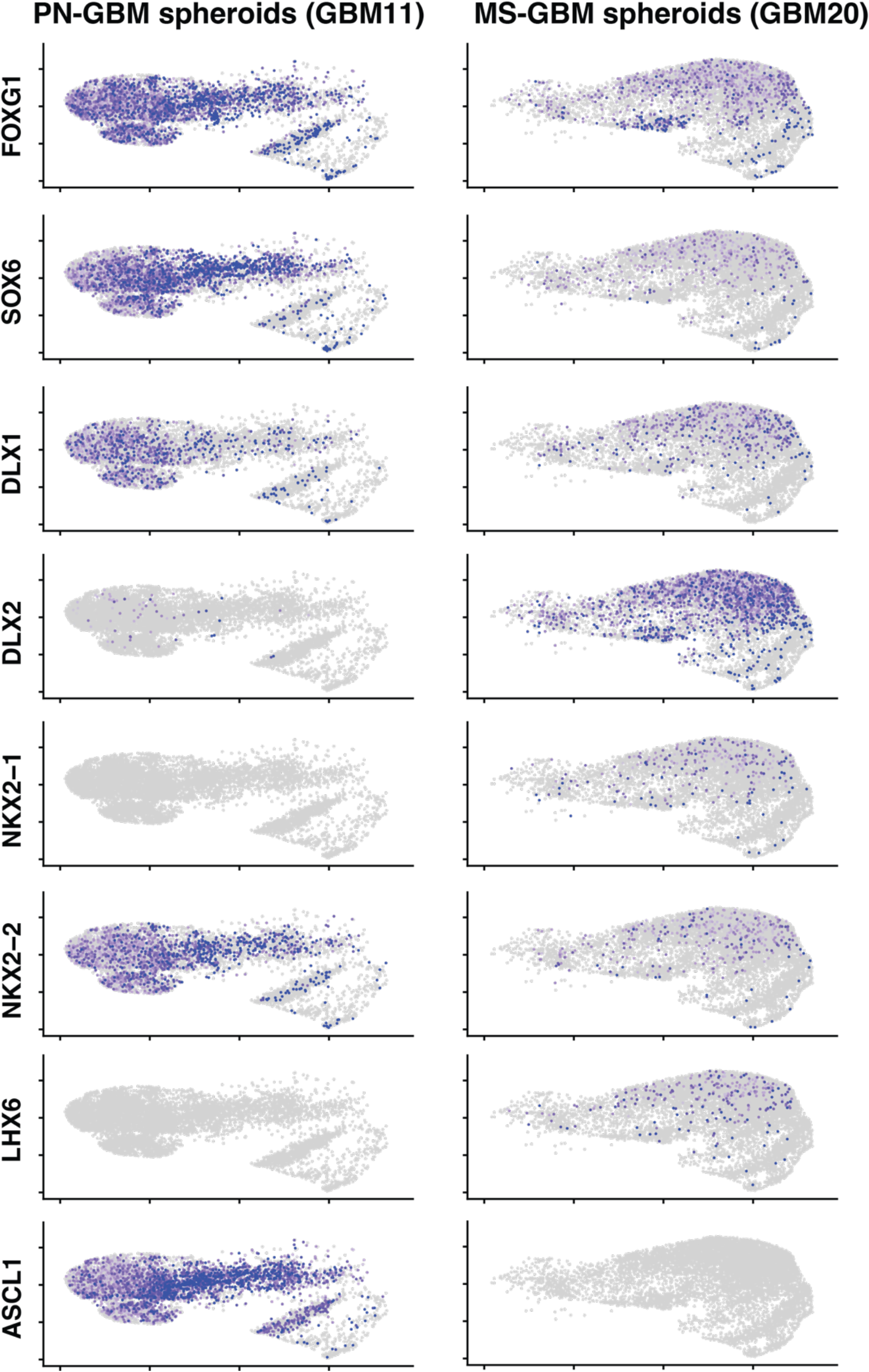
Feature plots of expression levels for canonical subpallial markers in PN (GBM11) and MS (GBM20) spheroids.

**Supplementary Figure 3.**
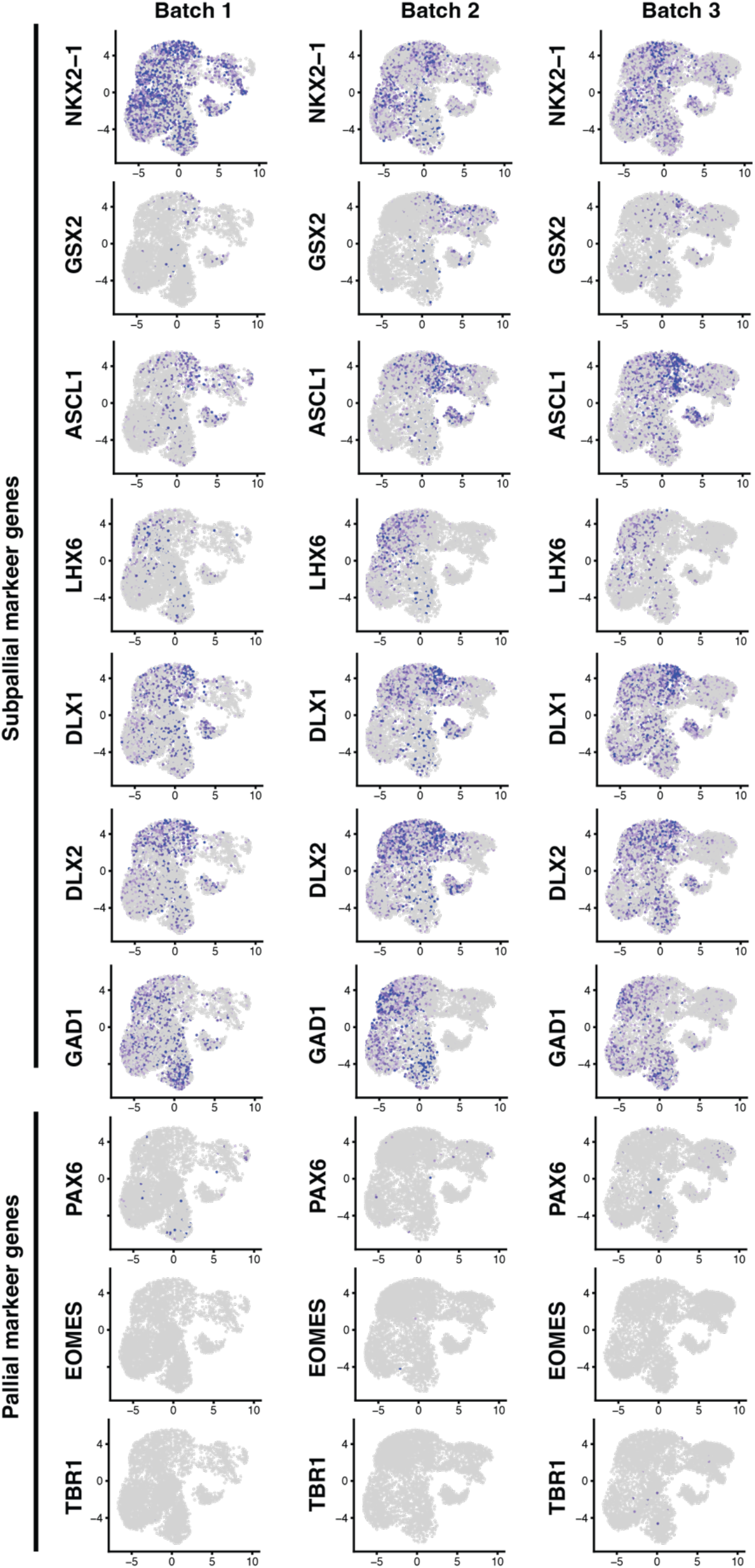
Feature plots of expression levels for pallial and subpallial marker genes in 3 independent batches of SNR-derived organoids.

**Supplementary Figure 4.**
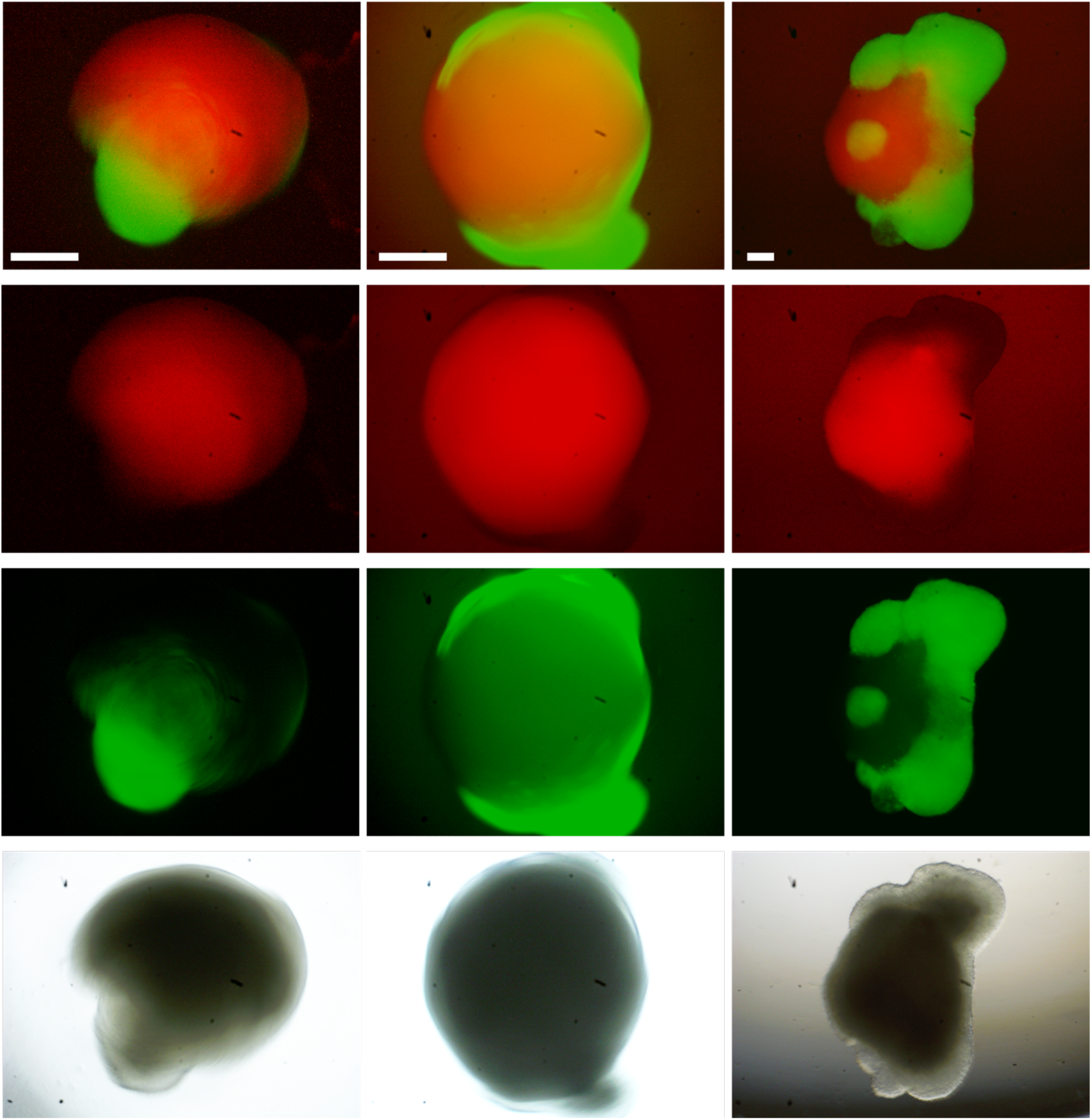
Images of iPSC-RFP-derived-organoids-MS-GBM chimeras. Fluorescent (organoid:red, GBM:green) images are on the top rows, and bright field images are on the bottom row.

**Supplementary Figure 5.**
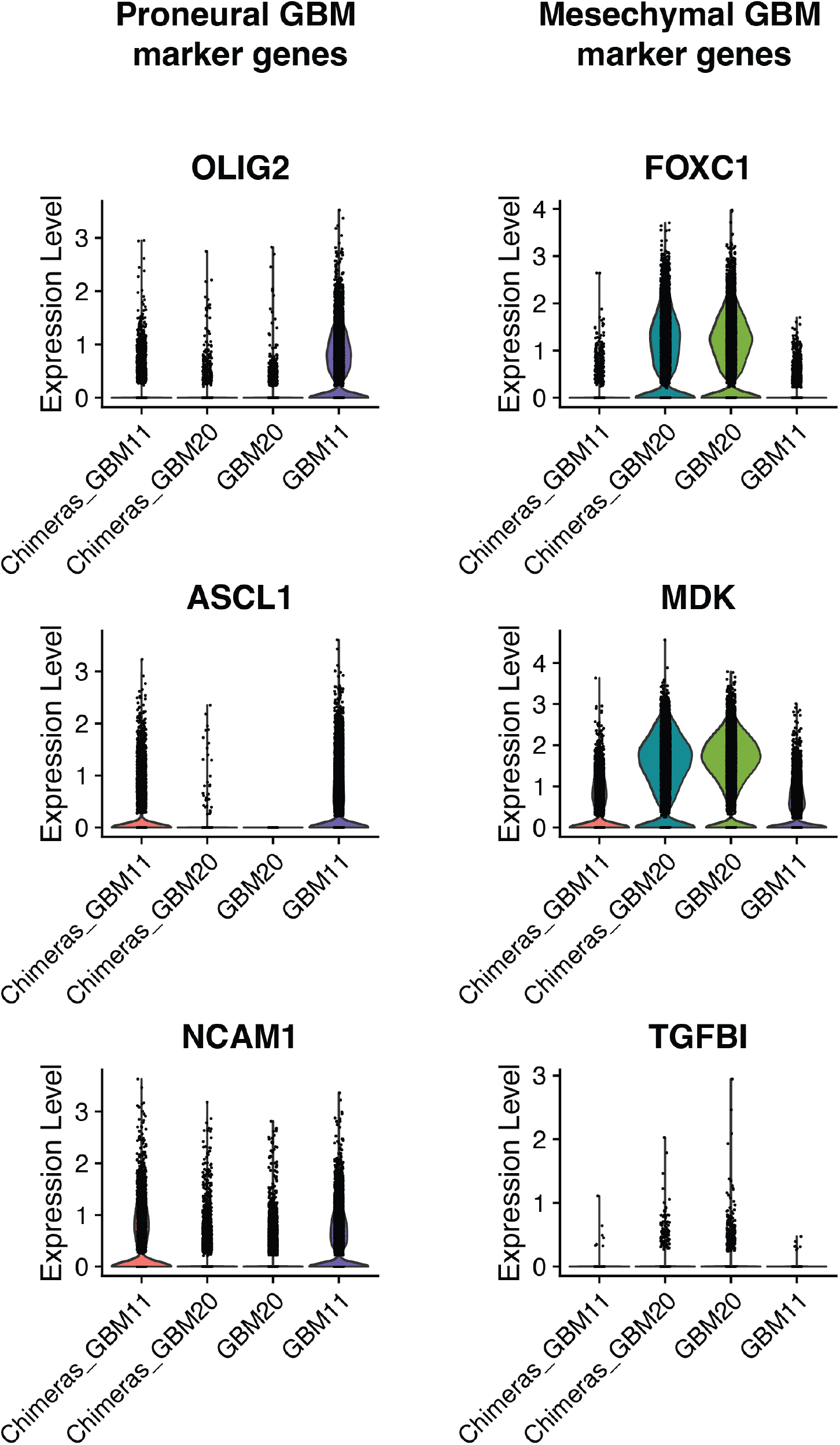
Violin plot of selected PN and MS markers in tumor cells from PN (GBM11) and MS (GBM20) spheroids and chimeras.

